# Methplotlib: analysis of modified nucleotides from nanopore sequencing

**DOI:** 10.1101/826107

**Authors:** Wouter De Coster, Mojca Strazisar

## Abstract

**Summary:** Modified nucleotides play a crucial role in gene expression regulation. Here we describe methplotlib, a tool developed for the visualization of modified nucleotides detected from Oxford Nanopore Technologies sequencing platforms, together with additional scripts for statistical analysis of allele specific modification within subjects and differential modification frequency across subjects.

**Availability and implementation:** The methplotlib command-line tool is written in Python3, is compatible with Linux, Mac OS and the MS Windows 10 Subsystem for Linux and released under the MIT license. The source code can be found at https://github.com/wdecoster/methplotlib and can be installed from PyPI and bioconda. Our repository includes test data and the tool is continuously tested at travis-ci.com.

## Introduction

Epigenetic nucleotide modifications, genome modifications that do not alter the primary DNA sequence, have many functions including transposon repression, gene expression regulation during development, imprinted gene expression and X-chromosome silencing (Gigante *et al.*, 2019; Greenberg and Bourc’his, 2019), and have been shown to play a role in many cellular functions, development, and pathological states such as psychiatric disorders and neurodegeneration (Gaine *et al.*, 2019; Armstrong *et al.*, 2019). Over 40 verified types of covalent nucleotide modifications have been described, of which 5-methylcytosine (5mC) and N6-methyladenine (m^6^A) are the most studied (Sood *et al.*, 2019). The long read sequencing platforms from Oxford Nanopore Technologies (ONT) enable genome-wide direct observation of modified nucleotides by assessing deviating current signals, for which multiple tools have been developed (McIntyre *et al.*, 2019; Liu, Fang, *et al.*, 2019; Liu, Georgieva, *et al.*, 2019; Rand *et al.*, 2017; Stoiber *et al.*, 2016; Simpson *et al.*, 2017), but a comprehensive evaluation of their performance is lacking. To the best of our knowledge, no flexible genome browser visualization method is available for this type of data.

## Methods

We developed *methplotlib*, a software package for visualization of the modified frequency and the per-read per-nucleotide probability of the presence of a nucleotide modification, together with additional summary overviews. While most work has been done on methylation, visualization using our tool is essentially agnostic to the type of nucleotide modification used as input, and future work may train upstream tools to recognize e.g. hydroxymethylation or various RNA modifications in direct RNA sequencing (Garalde *et al.*, 2018). At the time of writing no community-standard data format for nucleotide modifications has been established. The current methplotlib version is tailored to nanopolish (Simpson *et al.*, 2017), but the API can straightforwardly be expanded to accommodate modification data in CRAM/BAM, VCF, bigwig and custom formats from emerging tools, including direct modification identification during basecalling with the latest version of the Guppy basecaller. The input data is a tab-separated file from nanopolish containing either per read and per position likelihood of the presence of a modification, or the per position frequency of having a modified nucleotide across all reads. Gene and transcript annotation is extracted from a GTF file, and other types of annotation can be added in BED format.

Our methplotlib tool depends on core Python modules and numpy (Walt *et al.*, 2011), pandas (McKinney, 2011), scikit-learn (Pedregosa *et al.*, 2011), pyranges (Stovner and Sætrom, 2019) and plotly (Plotly Technologies Inc., 2015). We made our software easily available through PyPI and bioconda (Grüning *et al.*, 2018). Visualizations are created in dynamic HTML format, containing, optionally for multiple samples, i) the raw likelihood of nucleotide modification per position per read, ii) the frequency of having a modified nucleotide per position and iii) an annotation track, showing the exon and gene structure. The examples (Figure 1 and Supplementary Figures) were created using nanopolish call-methylation (Simpson *et al.*, 2017) of ONT PromethION data from a lymphoblastoid cell line of the Yoruban reference individual NA19240 (De Coster *et al.*, 2019) using annotation from Ensembl (Frankish *et al.*, 2019) and DNase hypersensitivity from ENCODE (ENCODE Project Consortium, 2012).

**Figure 1:**
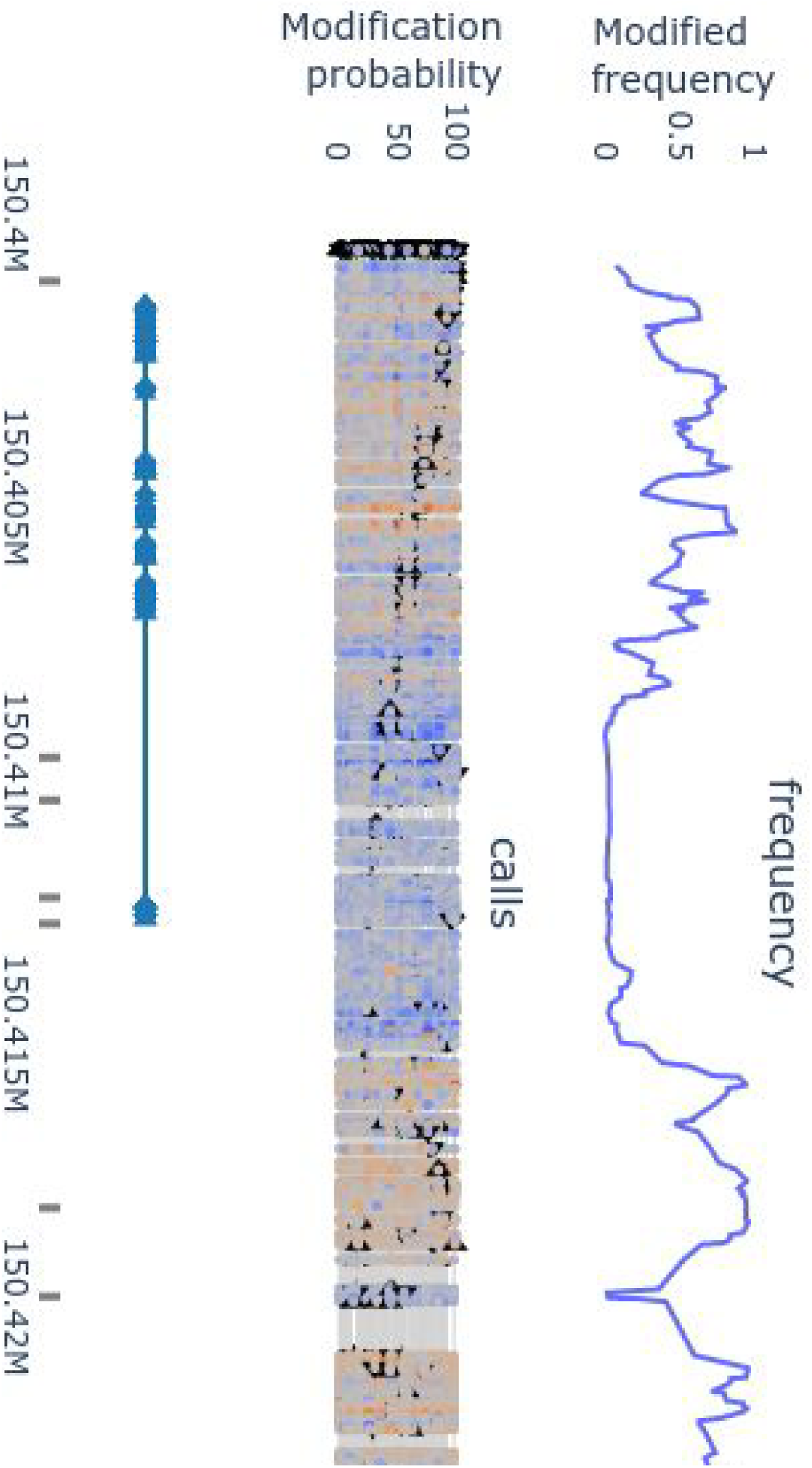
CpG methylation around the highly expressed CD74 gene. The upper panel shows per read per position the likelihood of a modified nucleotide with a color gradient (blue: low probability of modification, orange: high probability), the middle panel shows per position the frequency of methylated nucleotides, and the bottom panel contains gene structure annotation and DNase hypersensitive regions. Regulatory regions show lower frequency of methylation.

In addition, quality control plots are produced, including a principal component analysis to identify outliers, a pairwise correlation plot, highlighting more similar samples (Supplementary Figure S2), box plots of global modification frequencies and a bar chart of all positions for which modifications were identified.

Together with the tool we have also developed a snakemake workflow to facilitate processing of multiple datasets and multiple regions of interest (Koster and Rahmann, 2012). A companion script *annotate_calls_by_phase.py* is included to separate the modification results in both paternal haplotypes using a phased bam file from WhatsHap haplotag (Martin *et al.*, 2016). Using phased modification calls allows us to detect allele specific modification, statistically implemented using a Fisher exact test aggregating over a regulatory region (e.g. DNase hypersensitivity mark) in *allele_specific_modification.py*. This identifies mainly promoters affected by X-chromosome silencing (Supplementary Figure S3) and multiple known imprinted genes including *GNAS/GNAS-AS* (Supplementary Figure S4) (Weinstein *et al.*, 2010), *HYMAI1/PLAGL1* (Iglesias-Platas *et al.*, 2013) *and HERC3/NAP1L5* (Cowley *et al.*, 2012). In larger cohorts this approach could be used for identification of methylation quantitative trait loci. The same approach is straightforwardly expanded to differential modification testing in *differential_modification.py*, for example to test epigenetic differences between patients and unaffected subjects.

## Conclusion

Long read sequencing technologies of ONT and PacBio enable for the first time genome-wide direct observation of multiple types of nucleotide modifications without chemical modifications or affinity purification. To facilitate research in this emerging field we have developed methplotlib, a genome browser tool for the visualisation of per read raw nucleotide modification probabilities or aggregated frequencies derived from nanopore sequencing. Our package additionally includes a scalable workflow, quality control plots and scripts for statistical analysis. The API to import data can straightforwardly be expanded to use emerging data formats and multiple types of nucleotide modifications as identified by upstream software.

## References

Armstrong, M.J. et al. (2019) Diverse and Dynamic DNA Modifications in Brain and Diseases. Hum. Mol. Genet.

Cowley, M. et al. (2012) Epigenetic control of alternative mRNA processing at the imprinted Herc3/Nap1l5 locus. Nucleic Acids Res., 40, 8917–8926.

De Coster, W. et al. (2019) Structural variants identified by Oxford Nanopore PromethION sequencing of the human genome. Genome Res.

ENCODE Project Consortium (2012) An integrated encyclopedia of DNA elements in the human genome. Nature, 489, 57–74.

Frankish, A. et al. (2019) GENCODE reference annotation for the human and mouse genomes. Nucleic Acids Res., 47, D766–D773.

Gaine, M.E. et al. (2019) Differentially methylated regions in bipolar disorder and suicide. Am. J. Med. Genet. B Neuropsychiatr. Genet., 180, 496–507.

Garalde, D.R. et al. (2018) Highly parallel direct RNA sequencing on an array of nanopores. Nat. Methods, 15, 201–206.

Gigante, S. et al. (2019) Using long-read sequencing to detect imprinted DNA methylation. Nucleic Acids Res.

Greenberg, M.V.C. and Bourc’his, D. (2019) The diverse roles of DNA methylation in mammalian development and disease. Nat. Rev. Mol. Cell Biol., 20, 590–607.

Grüning, B. et al. (2018) Bioconda: sustainable and comprehensive software distribution for the life sciences. Nat. Methods, 15, 475–476.

Iglesias-Platas, I. et al. (2013) Imprinting at the PLAGL1 domain is contained within a 70-kb CTCF/cohesin-mediated non-allelic chromatin loop. Nucleic Acids Res., 41, 2171–2179.

Koster, J. and Rahmann, S. (2012) Snakemake--a scalable bioinformatics workflow engine. Bioinformatics, 28, 2520–2522.

Liu, Q., Fang, L., et al. (2019) Detection of DNA base modifications by deep recurrent neural network on Oxford Nanopore sequencing data. Nat. Commun., 10, 2449.

Liu, Q., Georgieva, D.C., et al. (2019) NanoMod: a computational tool to detect DNA modifications using Nanopore long-read sequencing data. BMC Genomics, 20, 78.

Martin, M. et al. (2016) WhatsHap: fast and accurate read-based phasing. bioRxiv, 085050.

McIntyre, A.B.R. et al. (2019) Single-molecule sequencing detection of N6-methyladenine in microbial reference materials. Nat. Commun., 10, 579.

McKinney, W. (2011) pandas: a foundational Python library for data analysis and statistics. Python for High Performance and Scientific Computing, 1–9.

Pedregosa, F. et al. (2011) Scikit-learn: Machine Learning in Python. J. Mach. Learn. Res., 12, 2825–2830.

Plotly Technologies Inc. (2015) Collaborative data science.

Rand, A.C. et al. (2017) Mapping DNA methylation with high-throughput nanopore sequencing. Nat. Methods, 14, 411–413.

Simpson, J.T. et al. (2017) Detecting DNA cytosine methylation using nanopore sequencing. Nat. Methods, 14, 407–410.

Sood, A.J. et al. (2019) DNAmod: the DNA modification database. J. Cheminform., 11, 30.

Stoiber, M.H. et al. (2016) De novo Identification of DNA Modifications Enabled by Genome-Guided Nanopore Signal Processing. bioRxiv, 094672.

Stovner, E.B. and Sætrom, P. (2019) PyRanges: efficient comparison of genomic intervals in Python. bioRxiv, 609396.

Walt, S. van der et al. (2011) The NumPy Array: A Structure for Efficient Numerical Computation. Comput. Sci. Eng., 13, 22–30.

Weinstein, L.S. et al. (2010) The role of GNAS and other imprinted genes in the development of obesity. Int. J. Obes., 34, 6–17.

