## Supplementary Figures for "Methplotlib: analysis of modified nucleotides from nanopore sequencing"

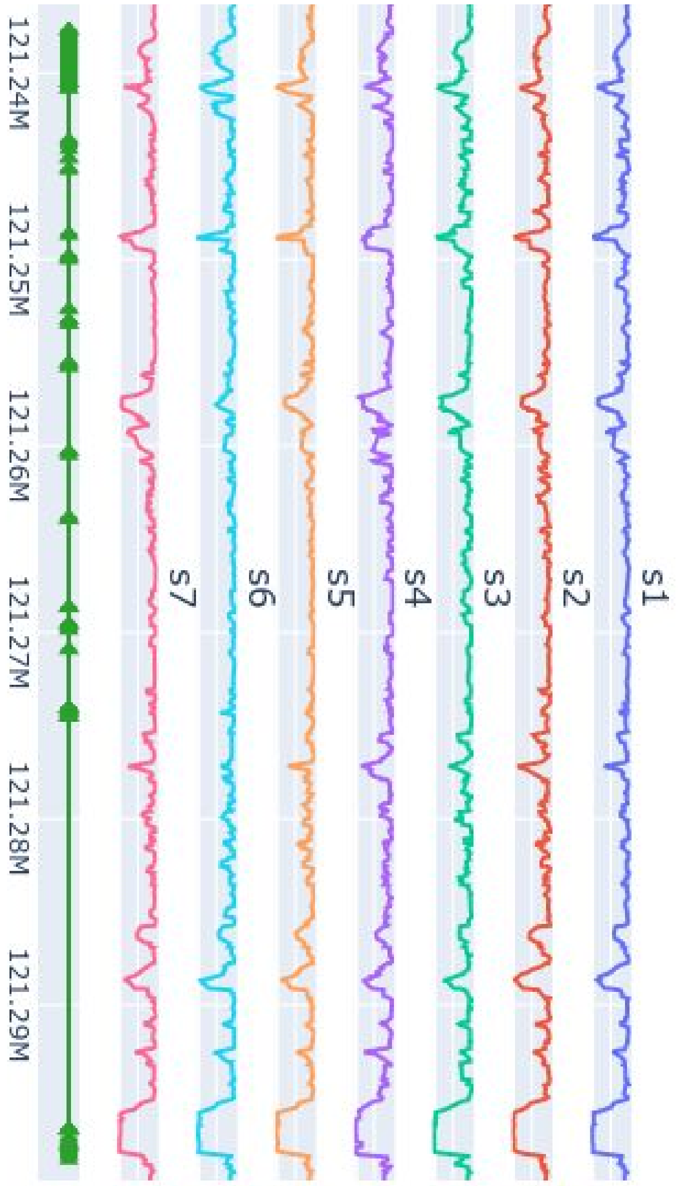


**Supplementary Figure S1: methylated frequency of multiple samples**
This plot shows the methylated frequency of 7 samples across the CAMKK2 gene.

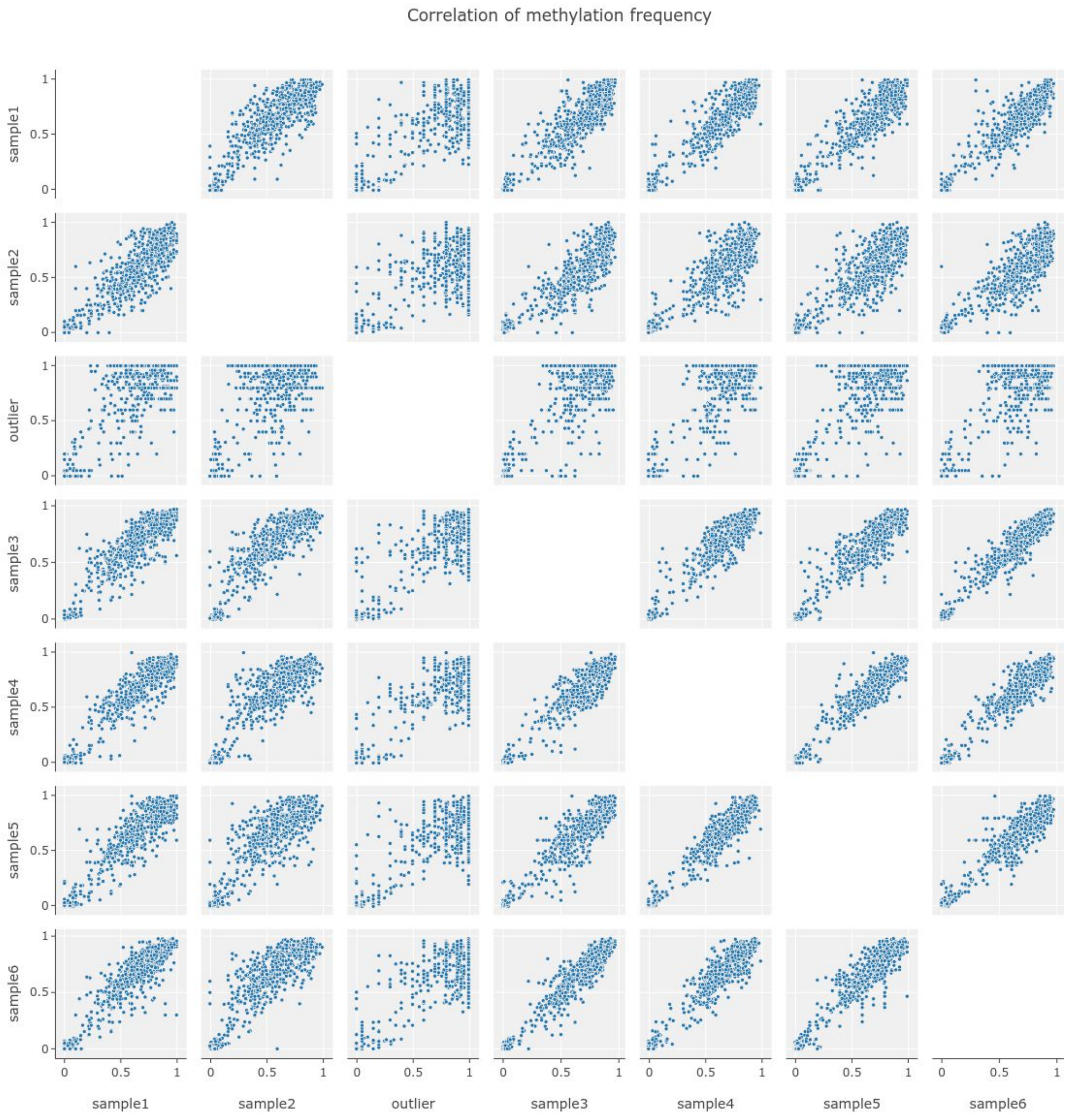


**Supplementary Figure S2: pairwise correlation plot**
Correlation of modification frequency across multiple datasets is plotted for each pair in the dataset, showing similarity for most samples with one notable outlier from a different tissue. All samples were sequenced on ONT PromethION.

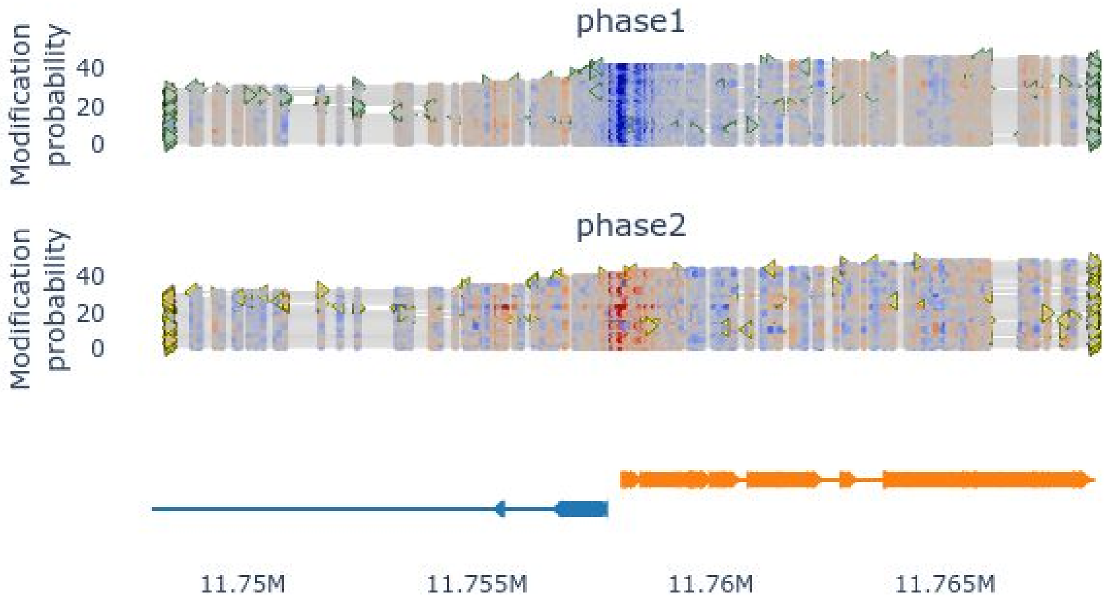


**Supplementary Figure S3: X-chromosome silencing**
Example of the MSL3 and AC004554.2 promoter, reads phased with WhatsHap

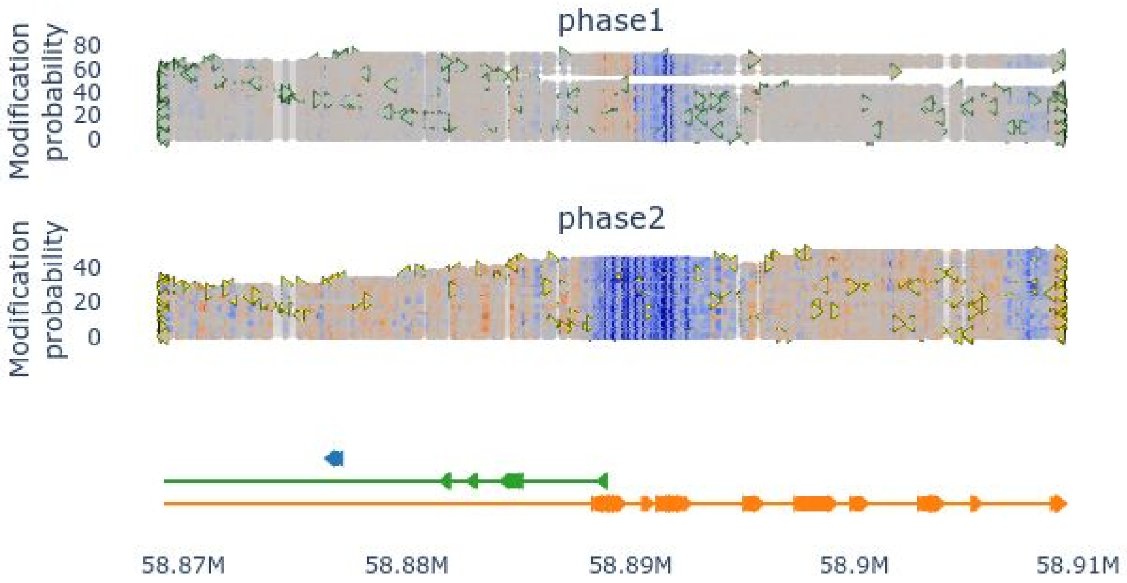


**Supplementary Figure S4: genomic imprinting**
Example of the GNAS/GNAS-AS locus, reads phased with WhatsHap
